# Chronic Δ9-Tetrahydrocannabinol Vapor Self-Administration Modulates Gut Microbiota and the Immune and Endocannabinoid Systems in Female Sprague- Dawley Rats

**DOI:** 10.64898/2026.08.30.748024

**Authors:** José Javier Rosado-Franco, Alysha L. Ellison, Elise M. Weerts, Catherine F. Moore, Dionna W. Williams

## Abstract

Vaping of cannabis and its primary components, including Δ9- tetrahydrocannabinol (THC), has increased during the last decade, especially among adolescents and adults. While the behavioral consequences of THC, including anxiety, depression, and associated impacts on reward-related signaling, are well documented, evaluation of additional pathways that have implications for neuronal function remains relatively unexplored. Further, despite evidence of the impact of THC on several physiologic processes, most studies have focused on the brain even though the endocannabinoid system with which THC interacts has widespread expression and functions throughout the body. To address these gaps, we obtained biological samples from adult female Sprague-Dawley rats (N=5-6 per group; all females) that were passively exposed to THC or vehicle vapor and then allowed to self-administer vaporized THC or vehicle intermittently for nine months. Animals were euthanized, and brain and peripheral organs were collected for analyses. We evaluated genes of endocannabinoid system-related receptors, modulators, ion channels, and metabolic enzymes, along with immune and microbiome-related pathways in 9 peripheral organs, 14 brain regions, and plasma. We observed that rats exposed to chronic intermittent THC vapor administration had widespread modulation of several endocannabinoid- system genes only in the brain, with most changes occurring in the dorsal striatum and hippocampus. In contrast, endocannabinoid-system gene expression in all evaluated peripheral organs remained unchanged after THC exposure. Similar findings occurred with immune-relevant genes, where *il-1β*, *il-6*, *ccl2*, *and mx1* were modulated only in the brain while expression in peripheral organs remained unchanged. Immune system modulation following chronic THC exposure significantly elevated plasma concentrations of IL-17A, gut microbiota dysbiosis, and global changes to circulating metabolites, including several amino acids, fatty acids, aminosugar, and nucleic acids. Our findings provide, for the first time, a whole-body evaluation of the molecular effects of chronic THC exposure throughout adulthood on the endocannabinoid and immune systems that have implications for cannabinoid-related behavioral effects.

## 1. Background

The use of cannabis and its constituents, specifically the major psychoactive phytocannabinoid Δ9-Tetrahydrocannabinol (THC), has been steadily increasing in the last decade, gaining popularity among adolescents and adults who use it recreationally or for medicinal purposes^1–4^. Inhalation of cannabis and THC-containing products is the most common route of administration, with specifically vaping of cannabis/THC by inhaling aerosols from a battery-powered heating device gaining in popularity. Vaping is considered a safer alternative to smoking combustible cigarettes by the general public, as it reduces exposure to tar and hundreds of toxic chemicals. However, it is important to note that vaping THC is not completely without risk, as it still produces carcinogens, respiratory toxicants, and teratogens, known to adversely impact health^5^. Cannabis and THC use can cause deleterious effects on behavior (i.e., development of anxiety, depression, dependence, etc.), cognition, immune system, aging, and accelerate the development of neurocognitive disorders^6–13^. Immediately following vapor inhalation, THC is rapidly absorbed into the blood and brain, producing higher parent drug concentrations and greater subjective effects compared with a similar dose administered by other routes^14–15^. However, a comprehensive molecular understanding of the mechanisms of harms caused by cannabis/THC inhalation, and their impact on THC-associated behavioral changes, remains incompletely characterized.

Psychiatric disorders have been broadly correlated with an inflammatory phenotype, including overexpression of pro-inflammatory cytokines, such as the IL-17 family, IL-6, and TNF-α, suggesting a skewed immune response toward a Th17 and regulatory T cell profile, and are often associated with gut microbiota dysbiosis, with lower abundance of bacteria that produce anti-inflammatory metabolites ^8,12–13,16–18^. While studies with human subjects are incredibly valuable, accessibility of organs, specifically the brain, is limited, that preclude comprehensive evaluation of the molecular effects of long-term use of THC across central and peripheral organ systems. To this end, our study aimed to bridge this gap by evaluating the impact of THC on endocannabinoid system receptors, modulators, and transporters and immune system genes across multiple organs, soluble cytokines, gut microbiota dysbiosis, and circulating metabolites.

In a study published previously, Moore *et al*. reported that female Sprague-Dawley rats chronically self-administered THC vapor^19–20^. In the present study, we used the brain and peripheral organs from a subset of these animals to evaluate the endocannabinoid and immune systems, gut microbiome, and circulating metabolite changes following THC or vehicle vapor exposure. We report that chronic THC exposure promoted substantial changes in endocannabinoid and immune system genes across several brain regions. However, to our surprise, these changes were restricted to the brain, as endocannabinoid and immune system genes remained unchanged in nine peripheral organs. These changes in brain gene expression were accompanied by significantly elevated circulating IL-17A, gut microbiota dysbiosis, and their circulating metabolic byproducts. Thus, our findings provide comprehensive evidence of physiological changes induced by chronic exposure to THC vapor that may be mechanistically linked to its behavioral effects, suggesting potential therapeutic pathways that may be effective in modulating adverse behavioral effects following THC vapor use.

## 2. Methods

### 2.1 Ethics Statement

All experimental protocols were approved by the Johns Hopkins University Animal Care and Use Committee. Animal handling, euthanasia, and biosample collection were conducted in accordance with the USDA Animal Welfare Regulations and the NIH Guide for the Care and Use of Laboratory Animals.

### 2.2 Animal subjects

Eleven female Sprague-Dawley rats (Charles River, MA, USA), approximately eight weeks old at the beginning of the experiment, were single-housed in wire-topped plastic cages in temperature- and humidity-controlled facilities with a reverse light cycle (12 hours, lights off from 8:00am-8:00pm). Rats were first exposed to passive vaporized administration of vehicle (100% propylene glycol; PG) or THC (200 mg/mL) intermittently (every other day) for two weeks. After a one-month wash-out period, rats were trained to self-administer puffs of either vehicle or THC (various concentrations: 5- 200 mg/mL). Self-administration sessions occurred every other day for eight months. Rats were provided standard corn-based chow (Teklad Diet 2018; IN, USA) to maintain at 90% free-feeding weight. Rats had *ad libitum* water access, except during experimental test sessions. Full methods details and behavioral results from this study were published previously and demonstrated that vaporized THC produced conditioned rewarding effects and supported self-administration behavior^19–20^.

Rats were approximately one year old at the end of the above study when biosamples were collected. One day (24 hours) following their last vapor self-administration session, animals were sedated with isoflurane and euthanized by rapid decapitation before necropsy. Upon necropsy, brain, blood, pancreas, heart, lung, kidneys, liver, spleen, skeletal muscle, adipose tissue, colon, and feces were collected and flash-frozen and stored at −80°C until further use. The brain was subdivided into 14 brain regions: dorsal striatum (DS), globus pallidus (GP), nucleus accumbens (NA), ventral pallidum (VP), cerebellum (C), frontal cortex (FC), hippocampus (H), hypothalamus (HY), substantia nigra (SN), ventral tegmental area (VTA), occipital cortex (OC), parietal cortex (PC), temporal cortex (TC), and thalamus (T). Necropsy was consistently performed at 8 AM to minimize circadian effects on endocannabinoid receptor expression.

### 2.3 RNA Extraction and cDNA Synthesis

RNA was extracted using the RNeasy Mini kit (Qiagen #74104, MD, USA) following the manufacturer’s instructions. Approximately 50mg of each tissue was added to 2mL tubes containing Lysing Matrix D (MP Biomedicals #11693050, CA, USA). Tissue was homogenized using MP FastPrep®-24 (MP Biomedicals #116913050-CF, CA, USA). The aqueous phase was mixed with 70% ethanol at a 1:1 ratio and loaded into the RNeasy columns. RNA-free DNase (Qiagen #79256, CA, USA) was added to the column to digest DNA present in the sample. RNA concentration and quality parameters were determined using Nanodrop (ThermoFisher Scientific, MA, USA). cDNA was synthesized using the iScript cDNA Synthesis kit (Bio-Rad #1708891, CA, USA) following the manufacturer’s instructions.

### 2.4 Relative Genetic Expression Determination

Quantitative Polymerase Chain Reaction (qPCR) was performed to determine the relative genetic expression of 17 genes from the immune and endocannabinoid systems using commercially available TaqMan primers as previously reported^21–22^.

### 2.5 Cytokine and Chemokine Determination

Cytokines and chemokines were quantified from rat plasma using the BioLegend LEGENDplex technology. Briefly, rat plasma was diluted 1:4, and the Rat Inflammation Panel (13-plex) V02 with Filter Plate (BioLegend, #741395; CA, USA) was used to determine the plasma concentration of cytokines and chemokines, according to the manufacturer’s protocol. Cytokine and Chemokine levels were determined using a BD FACSLyrics and analyzed using Qognit software version 2025-05-01.

### 2.6 Alpha and Beta Diversity and Taxonomic Changes Determination

DNA was extracted from feces samples using the QIAmp Powerfecal Pro DNA kit (Qiagen, #51804, MD, USA) following the manufacturer’s instructions. DNA quality parameters and concentration were determined using a Nanodrop (ThermoFisher Scientific, MA, USA). DNA was sent to Novogene (Sacramento, CA, USA) for library preparation, sequencing of the 16S Hypervariable Region 4, and bioinformatic analyses.

### 2.7 Biochemical Compound Quantification Using Metabolomics

Unbiased global metabolomics (ultra-high performance liquid chromatography and gas chromatography) was performed using plasma obtained at the time of necropsy. Samples were shipped to Metabolon© for quantification of circulating metabolites.

### 2.8 Data Analysis and Statistics

Determination of cytokines and chemokines, and relative gene expression, was done in duplicate and represented in graphs plotting the mean ± SD. For relative gene expression, our limit of detection (LoD) was calculated using an average of all animal subjects probing for Pan Eukaryotic 18S (ThermoFisher Scientific, #4333760F, MA, USA) with a cycle threshold of 35. Samples that did not amplify were given an arbitrary value of 39.99. Data were analyzed using PRISM software version 10.1 (GraphPad Software, Inc., San Diego, CA). Relative genetic expression variance between the relative expression of genes between organs was determined using Student’s t-test. Taxonomical variance was determined using Student’s t-test and LEfSE analyses. Biochemical quantification was determined using fold changes and p-values of the variance of these circulating metabolites.

## 3. Results

### 3.1: Endocannabinoid System Receptors, Modulators, Transporters, and Metabolic Enzymes are Primarily Upregulated in the Dorsal Striatum and Hippocampus After Chronic THC Vapor Self-administration

It is well established that THC interacts with many proteins in the canonical and extended endocannabinoid system, namely endocannabinoid receptors, modulators, and transporters. To evaluate the impact of THC on the endocannabinoid system, we performed qPCR on tissues obtained from rats that were trained to self-administer vapor puffs of either THC or propylene glycol vehicle for nine months. Surprisingly, we determined that eight endocannabinoid system genes, *cnr1, ppara, pparg, trpv1, trpv2, faah, naaa,* and *adora2a* (p-values=0.0095, 0.0190, 0.0048, 0.0095, 0.0381, 0.0381, 0.0095, 0.0381, respectively) were upregulated in the dorsal striatum, a vital component of the basal ganglia, following THC exposure as compared to vehicle (**Figure-1**). Similarly, eight genes were upregulated in the hippocampus: *cnr1, ppara, pparg, trpv1, faah, naaa, 5ht1a,* and *adora2a* (p-values=0.0043, 0.0043, 0.0043, 0.0043, 0.0043, 0.0043, 0.0087, 0.0087, respectively). From these ten endocannabinoid system genes, we found that only three had consistent upregulation among diverse brain regions following THC administration: *cnr1, pparg,* and *trpv1*. In contrast, *cnr2* mRNA was downregulated by chronic THC exposure across multiple brain regions. The remaining endocannabinoid genes were modulated following THC exposure in a brain region-specific manner, and there were no consistent patterns of changes to their expression. We also evaluated *gpr18*, *gpr55*, *gpr110,* and *gpr119*, but found almost no detectable expression in any of the samples (data not shown), consistent with our previous findings^21–22^. We evaluated these endocannabinoid system genes in nine peripheral organs of these same rats and were surprised to find no changes relative to vehicle **(Supplemental Figure-1)**, which is in sharp contrast to the brain. These findings suggest brain-specific mechanisms occur following chronic THC exposure that are not conserved in peripheral organs.

**Figure-1:**
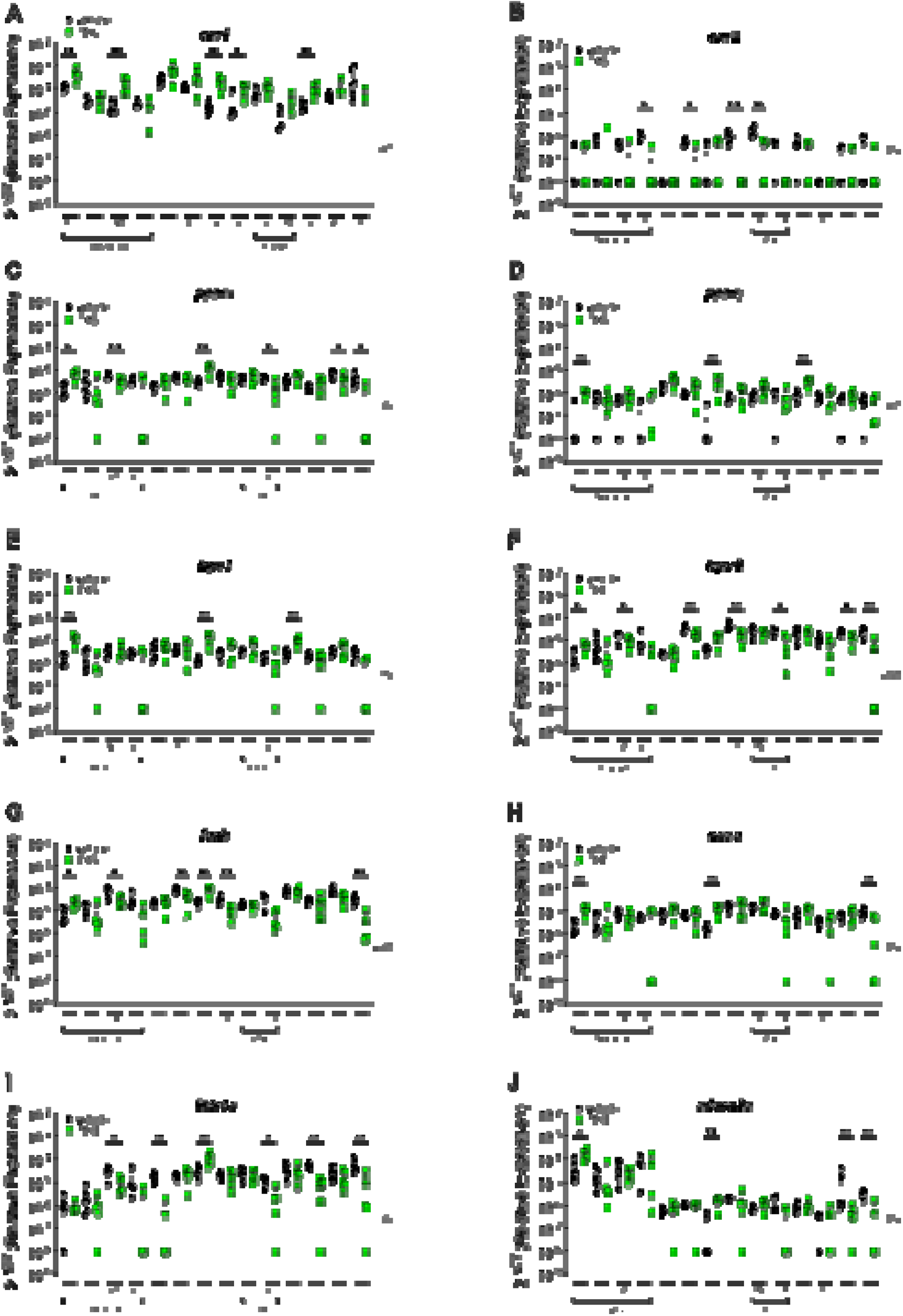
Endocannabinoid System Receptors, Modulators, Transporters, and Metabolic Enzymes are Mostly Upregulated in the Dorsal Striatum and Hippocampus After Chronic Self-administration of THC. Relative expression of **(A)** cnr1*, (**B)** cnr2, **(C)** ppara, **(D**) pparg, **(E)** trpv1, **(F)** trpv2, **(G)** faah,* **(H)** *naaa,* **(I)** *5ht1a (J) adora2a* was determined using qPCR from brain of female Sprague Dawley rats that received either vaped vehicle (black circles, n=6) or THC (green squares, n=5) in alternating days for eight months. DS=dorsal striatum, GP=globus pallidus, NA=nucleus accumbens, VP=ventral pallidum, C=cerebellum, FC=frontal cortex, H=hippocampus, HY=hypothalamus, SN=substantia nigra, VTA=ventral tegmental area, OC=occipital cortex, PC=parietal cortex, TC=temporal cortex, T=thalamus. Data were analyzed using one-way ANOVA and are represented using the mean ± SD.

### 3.2: Chronic THC Vapor Self-Administration Selectively Alters *il1b, il6, ccl2, mx1*, and *trail*, but not Type I Interferons (*ifna* and *ifnb*), in a Brain Region-Specific Manner

The interaction of THC and the endocannabinoid system is established to impact immune system pathways and hence, immune homeostasis. This has been primarily studied in a disease or infectious context. However, little is known about the impact of chronic THC administration on the immune system in the absence of disease. Therefore, we evaluated genes broadly implicated in immune response to evaluate whether they were modulated following THC exposure. From our selected immune markers, we found that *il-1*β*, il-6, ccl2, mx1* and *trail* had differential expression among many subregions of the brain (**Figure-2**). Surprisingly, we found that *il-1*β was significantly increased following THC exposure in the hippocampus (p-value=0.0152), *il- 6* was decreased in the cerebellum, hypothalamus and occipital cortex (p-values=0.0303, 0.0043, 0.303 respectively), *ccl2* was decreased in the nucleus accumbens, cerebellum and thalamus (p-values=0.0043, 0.303, 0.0043, respectively), while increased by THC in the ventral tegmental area (p-value=0.0159), *mx1* was significantly increased in the cerebellum, hippocampus and occipital cortex (p-values=0.0173, 0.0087, 0.0281, respectively), and *trail* was significantly increased by THC in the dorsal striatum, hippocampus and occipital cortex (p-values=0.0048, 0.0022, 0.0152 respectively). While THC exposure promoted broad changes in immune genes across multiple brain regions, there was a selective response to its effects, as *ifna* and *ifnb,* important antiviral markers, remained unchanged. Similar to the endocannabinoid system genes, the effects of THC exposure were restricted to the brain, as there were no changes in any of these genes among the nine peripheral organs evaluated **(Supplemental Figure-2).** Altogether, these results suggest that chronic THC exposure exerts pro-inflammatory effects, while maintaining antiviral capacity, in a brain sub- region-specific manner that warrants further investigation of the impact of THC use on neurologic immunity.

**Figure-2:**
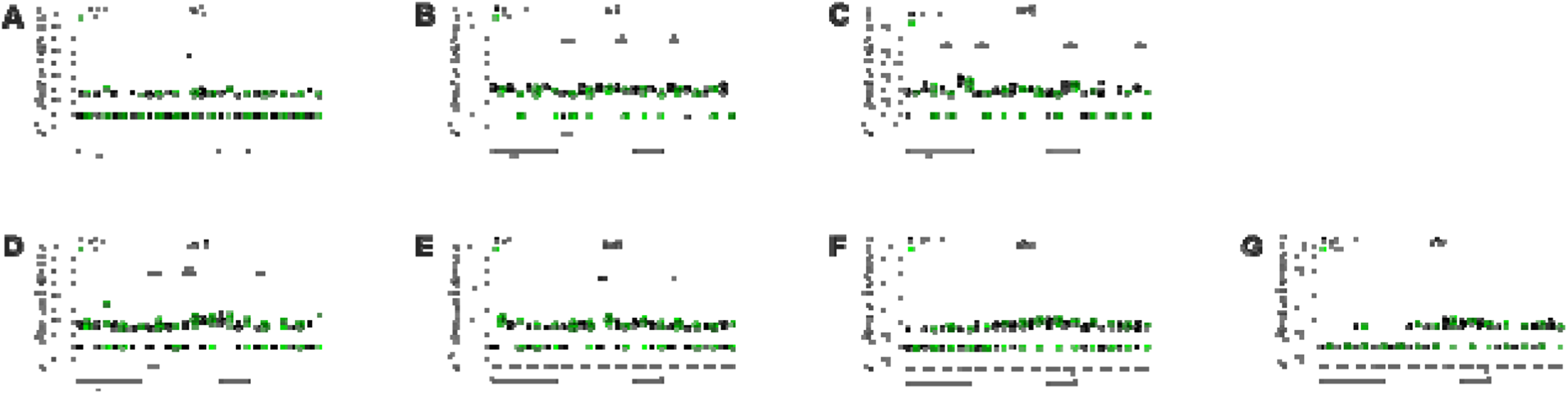
THC Chronic Self-Administration Selectively Alters *il1b, il6, ccl2, mx1*, and *trail*, but not Type I Interferons (*ifna* and *ifnb*), in a Brain Sub-region Manner. Relative expression of **(A)** *il1b,* **(B)** *il6,* **(C)** *ccl2,* **(D)** *mx1,* **(E)** *trail,* **(F)** *ifna, and* **(G)** *ifnb* was determined using qPCR from brain of female Sprague-Dawley rats that received either vaped vehicle (black circles, n=6) or THC (green squares, n=5) on alternating days for nine months. DS=dorsal striatum, GP=globus pallidus, NA=nucleus accumbens, VP=ventral pallidum, C=cerebellum, FC=frontal cortex, H=hippocampus, HY=hypothalamus, SN=substantia nigra, VTA=ventral tegmental area, OC=occipital cortex, PC=parietal cortex, TC=temporal cortex, T=thalamus. Data were analyzed using one-way ANOVA and are represented using the mean ± SD.

### 3.3: Chronic THC Vapor Self-administration Increases Plasma IL-17A

As we found a striking and selective modulation of immune genes in the brain, but not peripheral organs, we next evaluated plasma cytokines as they can readily cross the blood-brain barrier and impact brain function. We evaluated 13 cytokines and chemokines commonly implicated in immune regulation. Of the 13 measured analytes, we found that only IL-17A (p-value=0.0397) was increased in the animals who self-administered THC, compared to vehicle (**Figure-3**). Indeed, there was no significant effect of chronic THC exposure on any of the other evaluated cytokines, further indicating a well-controlled and regulated effect of the phytocannabinoid on the immune system under basal conditions in the absence of disease.

**Figure-3:**
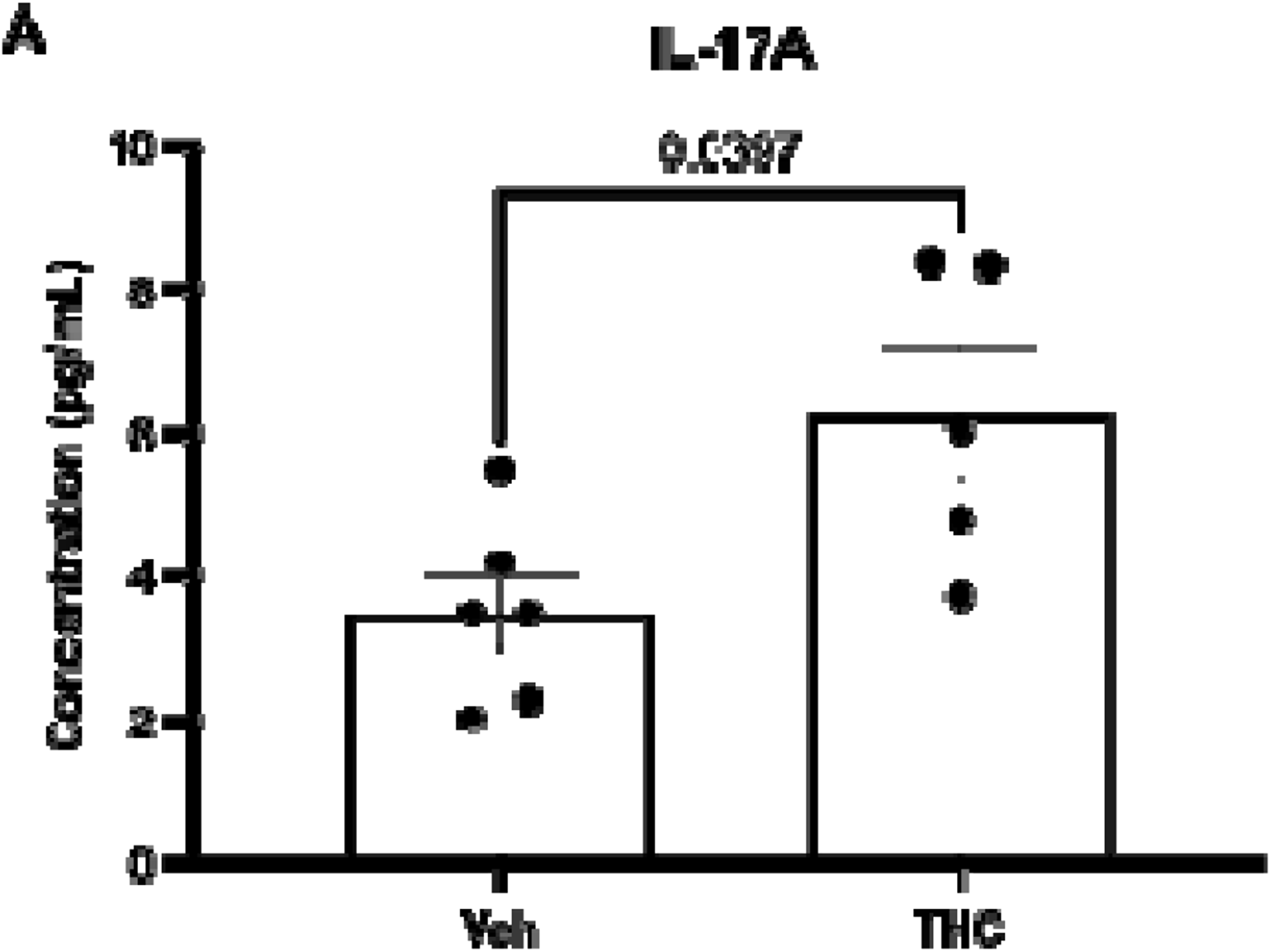
Chronic THC Self-administration Increases IL-17A levels. Serum samples were analyzed using LEGENDplex technology. **A)** Serum IL-17 concentrations from female Sprague-Dawley rats that received either vaped vehicle or THC. Data were analyzed using Student’s t-test and are represented using the mean ± SD.

### 3.4: Chronic THC Vapor Self-administration Increases Phylum Bacteroidetes Abundance but Reduces Firmicute Abundance

The microbiome is regulated by lifestyle factors and greatly impacts human health. Further, there is a direct relationship between microbiome composition, immunity, and brain functions. Thus, we next characterized gut microbiota composition following chronic THC vapor exposure. We found no differences in microbiome richness, evenness, alpha, or beta diversity (**Supplemental Figure-3**). However, changes in taxonomical composition occurred between the rats who self-administered THC vapor relative to vehicle. Rats who self-administered THC had microbiomes enriched with members of the phylum Bacteroidetes, in contrast to rats exposed to vehicle vapor that were enriched with Firmicutes (**Figure-4**). Moreover, members of the genus *Muribaculaceae* were enriched following THC self-administration (**Figure-4**). These results demonstrate that chronic THC vapor administration modulates gut microbiota, which is increasingly recognized for its role in psychiatric, immune, and energy metabolism-related disorders, which may be one potential mechanism by which chronic THC use promotes broad physiologic responses.

**Figure-4:**
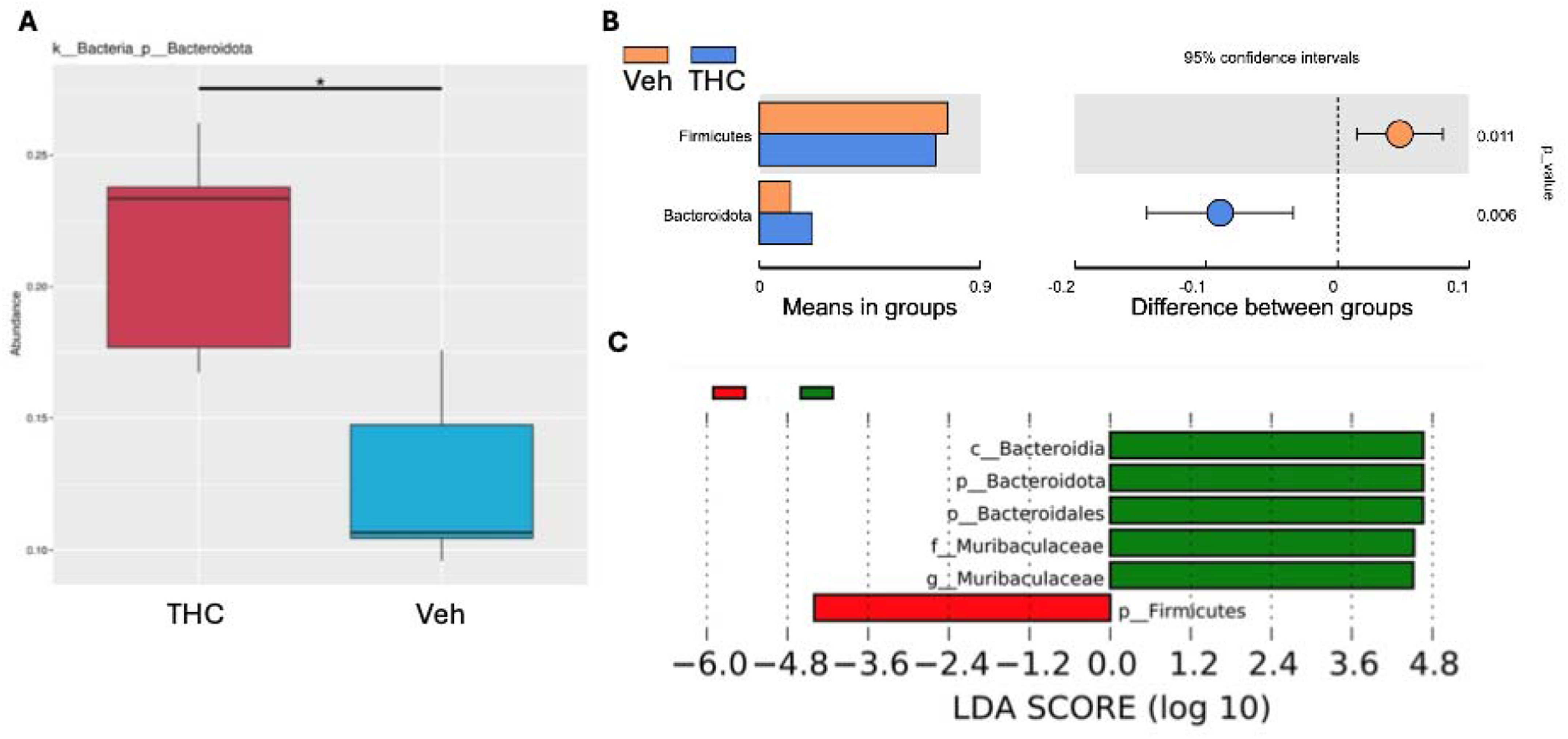
THC Chronic Self-administration Increases Phylum Bacteroidota Abundance, but Reduces Abundance of Firmicutes. DNA was extracted from the feces of female Sprague-Dawley rats who underwent chronic THC self-administration. **A)** Bacterial abundance in rats who self-administered THC (red square) vs vehicle (blue square). **B)** Student t-test results from Bacteroidetes and Firmicutes. **C)** LefSe analysis showing the taxonomical difference between the Vehicle group vs THC. Metastast analyses, Students’ t-test and LEfSe analyses were used to analyze this data.

### 3.5 Chronic THC Vapor Self-administration Alters Amino Acid, Creatine, Fatty Acid, Aminosugar, and Nucleic Acid Metabolism

As the gut microbiota contributes to the availability of circulating metabolites important for homeostasis, we performed unbiased metabolomics to evaluate whether there were functional changes to circulating metabolites following chronic THC self-administration. We identified 17 metabolites that were significantly regulated between rats who self-administered THC compared to vehicle. Of these 17 metabolites, 13 were increased in the rats that self-administered THC, while the remainder were decreased (**Table-1**). We found that chronic THC self-administration increased circulating metabolites involved in amino acid metabolism, such as glutamate, tyrosine, leucine, isoleucine, valine, methionine, and cysteine metabolism, S-adenosylmethionine & taurine metabolism, creatine, fatty acid, lysophospholipid, plasmalogen, and one plant component. In contrast, THC exposure decreased metabolites related to histidine, aminosugar, phospholipid, and purine metabolism (**Table-1**). Importantly, we found that glutamate metabolism was increased following THC exposure, indicated by the presence of Carboxyethyl-GABA (CEGABA) (fold change 1.26; p-value=0.0057), a derivative of the inhibitory neurotransmitter GABA. These results demonstrate that gut microbiota modulation that occurred following chronic THC exposure affected the availability of circulating metabolites that play crucial roles in maintaining brain and immune system homeostasis. Together, these findings suggest microbiome-immune-endocannabinoid pathways modulated following THC exposure as potential mechanisms by which chronic use contributes to psychiatric disorders, immune dysregulation, and altered metabolic processes.

**Table-1:** Chronic THC Self-administration Alters Amino Acid, Creatine, Fatty Acid, Aminosugar, and Nucleic Acid Metabolism. Serum samples were sent to the Emory Integrated Metabolomics and Lipidomics Core to determine the presence and differences in the fold change of biochemical compounds. Carboxyethyl GABA, vanillactate, Isovalerate, 2-hydroxy-3-methylvalerate, S-methylcysteine, guanidinoacetate, pentadecanoylcarnitine, linoleycholine, and dihydroferulic acid sulfate were upregulated, while Phosphoethanolamine, Urate, N-glycolylneuraminate, and N- acetylcarnosine were downregulated. Data are represented in significant fold changes and p-values.

| Increased Circulating Metabolites |  |  |  |
| --- | --- | --- | --- |
| Metabolite | Metabolic Pathway | Fold Change | p-value |
| Carboxyethyl-GABA<br>(CEGABA) | Glutamate metabolism | 1.26 | 0.0057 |
| Vanillactate | Tyrosine metabolism | 1.41 | 0.0435 |
| Isovalerate (i5:0) | Leucine, Isoleucine & Valine metabolism | 1.30 | 0.0118 |
| 2-hydroxy-3-methylvalerate | Leucine, Isoleucine & Valine metabolism | 1.76 | 0.0170 |
| S-methyl cysteine | Methionine, Cysteine, SAM & Taurine metabolism | 1.28 | 0.0301 |
| Guanidinoacetate | Creatine metabolism | 1.30 | 0.0187 |
| Pentadecanoylcarnitine (C15)* | Fatty acid metabolism | 1.49 | 0.0339 |
| Linoleoylcholine* | Fatty acid metabolism | 2.48 | 0.0373 |
| 1-palmitoleoyl-GPC (16:1)* | Lysophospholipid metabolism | 1.15 | 0.0488 |
| 1-tetracosadienoyl-GPC (24:2)* | Lysophospholipid metabolism | 1.25 | 0.0477 |
| 1-(1-enyl-palmitoyl)-2-palmitoleoyl-GPC (P-16:0/16:1)* | Plasmalogen metabolism | 1.57 | 0.0078 |
| Dihydroferulic acid sulfate | Food/Plant component | 2.93 | 0.0327 |
| **Decreased Circulating Metabolites** |  |  |  |
| **Metabolite** | **Metabolic Pathway** | **Fold Change** | **p-value** |
| N-acetyl carnosine | Histidine Metabolism | 0.80 | 0.0362 |
| N-glycolylneuraminate | Aminosugar metabolism | 0.75 | 0.0499 |
| Phosphoethanolamine | Phospholipid metabolism | 0.70 | 0.0069 |
| Urate | Purine metabolism | 0.76 | 0.0475 |

## 4. Discussion

Using a multi-systems approach, we report the effects of chronic THC vapor exposure by self-administration in adult female Sprague-Dawley rats, focusing on changes in endocannabinoid and immune gene expression, circulating cytokines and metabolites, and microbiome composition. Interestingly, THC modulated several endocannabinoid and immune genes that were restricted to the brain (especially the dorsal striatum and hippocampus) and remained unchanged across several peripheral organs. Further, these changes were accompanied by increased circulating IL-17A, changes in gut microbiota composition, and modulation of several metabolites, including those important for neurotransmission. Together, these processes collectively impact brain and physiologic health and demonstrate that chronic THC use is not benign. Importantly, our study occurred in the absence of infection or disease and is a starting point to begin understanding the long-term consequences of THC exposure in otherwise healthy animals with chronic use throughout adulthood. Altogether, our current findings identify fundamental mechanisms regarding chronic THC and its effect on endocannabinoid signaling, immune homeostasis, and the gut-brain axis.

Our model provides advantages over human studies, due to the experimental control facilitated by preclinical animal models that are nearly impossible to control in a human population (i.e., THC dose, frequency of use, reason for use, concomitant use of additional addictive substances, diet, and lifestyle); hence, our findings close the gap in knowledge regarding the molecular contributions of chronic THC use throughout adulthood and its impact in overall health.

The endocannabinoid and immune systems have co-evolved to ensure survival^8, 24–25^. Dysregulation of the immune system is increasingly recognized as a contributor to psychiatric conditions. Indeed, high levels of pro-inflammatory cytokines, such as IL- 17A, and gut microbiota dysbiosis are associated with psychiatric disorders^12,16–18,23–24^. In this context, our findings identify an important cellular target involved in chronic THC use: γδCD4+ T cells. Unlike other cytokines that are highly pleiotropic, IL-17A is a pro- inflammatory cytokine selectively produced by γδCD4+ T cells and by group 3 innate lymphoid cells to modulate Th17-effector functions^8^. Interestingly, the possibility exists that the THC-induced changes in circulating IL-17A were related to its effects on microbiome composition, as molecular signals from pathogenic or commensal bacteria have been shown to regulate IL-17A transcription^8^. Our microbiome findings are of additional importance as the *Muribaculaceae* genus maintains host health by producing short-chain fatty acids, degrading mucin, and regulating intestinal permeability and barrier integrity^26^. They also produce B vitamins for the host and metabolize numerous amino acids, such as aspartate and glutamate, known to have anxiogenic properties, as well as glutamine, glycine, methionine, valine, isoleucine, and leucine^26^. Thus, circulating metabolites altered by chronic THC use may directly impact mental health through changes to the gut microbiome by consequent effects on neuroendocrine and/or neuroactive compounds, and neurotransmitter regulation through vagus and mesenteric nerve stimulation^26^.

Our endocannabinoid system findings shed insight on coordinated and tightly regulated differential gene expression among brain subregions, identifying convergence between the dorsal striatum and the hippocampus - brain regions associated with motor control, action selection, and the formation of habits, and learning, memory, spatial navigation, and plasticity, respectively^28–33^. While most preclinical THC studies have primarily, and appropriately, focused solely on dopaminergic and reward-related pathways, we performed a comprehensive evaluation across several brain regions. To this end, we found eight endocannabinoid system genes upregulated in the dorsal striatum (*cnr1, ppara, pparg, trpv1, trv2, faah, naaa*, and *adora2a*) and hippocampus (*cnr1, ppara, pparg, trpv1, faah, naaa, 5htia,* and *adora2a*) after chronic THC use. Of importance, upregulation of *cnr1, ppara, pparg, trpv1, faah, naaa,* and *adora2a* was observed in both of these brain subregions. These findings demonstrate the importance of broadly evaluating the effects of THC across multiple brain subregions, as there may be unanticipated effects in understudied areas that may shed important insights into addiction, drug-seeking behavior, and reward cues/memory. It is important to acknowledge that our findings contrast with well-established downregulation of endocannabinoid tone conventionally known to occur following prolonged THC use, but these differences may be due to differences in the route of administration used, THC dose, biological sex, microbiome, and duration of THC exposure^37–38^.

Similar to our endocannabinoid system findings, we identified modulation of immune system genes by THC selectively in the brain, that did not occur in peripheral organs, highlighting the profound immunomodulatory effect of THC in the CNS. Importantly, pro-inflammatory cytokines, including IL-17A, are implicated in inflammatory pathways that promote tryptophan degradation and reduce serotonin availability, which may be an important aspect of THC-related behavioral effects^39^. While these findings did not reach statistical significance, we identified important trends in the tryptophan and serotonin metabolism following chronic THC exposure that warrant further investigation (**Supplementary Table-1**).

We report that chronic THC exposure altered several metabolic pathways that regulate neurotransmitters, anti-inflammatory, antioxidant, and antidiabetic compounds, as well as important components of energy metabolism, metabolic homeostasis, and cell membrane composition. Of utmost importance, our results identified an increase in CEGABA, a precursor of GABA. Previous results show that imbalances in GABA are caused by microbiota dysbiosis that can lead to anxiety and other neurocognitive disorders^34–36^. Furthermore, our identified increased metabolites have anti-inflammatory properties, including isovalerate, S-methyl cysteine, pentadecanoyl carnitine, and dihydroferulic acid. Additionally, metabolites decreased following THC use included N-acetyl carnosine, involved in histidine metabolism, preventing oxidative stress, inflammation, aging, and improving neuroprotection. Similarly, palmitoylethanolamide is related to phospholipid composition and mitochondrial structure and function, has antioxidant properties, neuroprotective effects, and helps maintain normal blood pressure levels, highlighting deleterious consequences of chronic THC use. These findings add to the existing literature regarding the adverse metabolic consequences of THC in lipid membrane metabolism, DNA/RNA pathways, and oxidative stress responses^5^.

We recognize that our study has the following limitations. We only included female Sprague-Dawley rats in this study; future work examining sex differences is of high importance. There were no longitudinal blood or feces samples to track the effect of acute THC use; only terminal samples were used. The route of administration, vaping, can not be generalized to other routes of administration used in other studies given known differences in THC pharmacokinetics and metabolic pathways. It is important to note that control animals had previous exposure to THC vapor, though this was a limited amount of exposure that occurred eight months prior to brain and organ collection^19^.

## 5. Conclusions

Together, our findings identify interconnected pathways involving endocannabinoid, immune, and microbiome systems that occurred following chronic THC use in adulthood. To the best of our knowledge, this is the first preclinical study demonstrating that chronic THC exposure in female rats alters brain endocannabinoid and immune system genes, increases IL-17A, and induces gut microbiota dysbiosis, effectively altering circulating metabolites known to maintain immune homeostasis and overall health.

## 6. List of Abbreviations

THC: Δ9-tetrahydrocannabinol
5htia: 5-hydroxytryptamine receptor 1A
*adora2a*: Adenosine A2A Receptor
CCL2: C-C motif chemokine ligand 2
*cnr1*: Cannabinoid Receptor 1
*cnr2*: Cannabinoid Receptor 2
CEGABA: Carboxyethyl aminobutyric acid
C: Cerebellum
DS: Dorsal striatum
GABA: Gamma-aminobutyric acid
GP: Globus pallidus
*faah*: Fatty Acid Amide Hydrolase
FC: Frontal cortex
H: Hippocampus
HY: Hypothalamus
IL-1β: Interleukin 1β
IL-17: Interleukin 17
IL-6: Interleukin 6
LoD: Limit of detection
*mxa*: Myxovirus resistance protein 1
*naaa*: N-Acylethanolamine Acid Amidase
NA: Nucleus accumbens
OC: Occipital cortex
PC: Parietal cortex
*ppara*: Peroxisome Proliferator-Activated Receptor Alpha
*(pparg*: Peroxisome Proliferator-Activated Receptor Gamma
qPCR: Quantitative Polymerase Chain Reaction
SN: Substantia nigra
TC: Temporal cortex
T: Thalamus
*trpv1*: Transient Receptor Potential Vanilloid 1
*trpv2*: Transient Receptor Potential Vanilloid 2
TNF-α: Tumor necrosis factor- α
VP: Ventral pallidum
VTA: Ventral tegmental area

## 7. Declarations

### 7.1 Ethics approval and consent to participate

Included in Methods, section 2.1 Ethics statement.

### 7.2 Consent for publication

Not applicable

### 7.3 Availability of Data and Materials

All data generated or analyzed during this study are included in this published article and its supplementary information file. The datasets used and/or analyzed during the current study are available from the corresponding author on reasonable request.

### 7.6 Author Contributions

Conceptualization: DWW

Methodology: ALE, JJRF, and CFM

Investigation: ALE, JJRF, and CFM

Funding acquisition: DWW

Project administration: DWW, CFM, and EW

Supervision: DWW

Writing – original draft: JJRF and DWW

Writing – review & editing: All authors

## Acknowledgments

We thank all members of the Williams Lab and the Department of Pharmacology and Chemical Biology at Emory University for their thoughtful conversations and input regarding the findings of this manuscript.

