## Supplementary figures and images for "Chronic Δ9-Tetrahydrocannabinol Vapor Self-Administration Modulates Gut Microbiota and the Immune and Endocannabinoid Systems in Female Sprague- Dawley Rats"

### Supplemental Figure-1

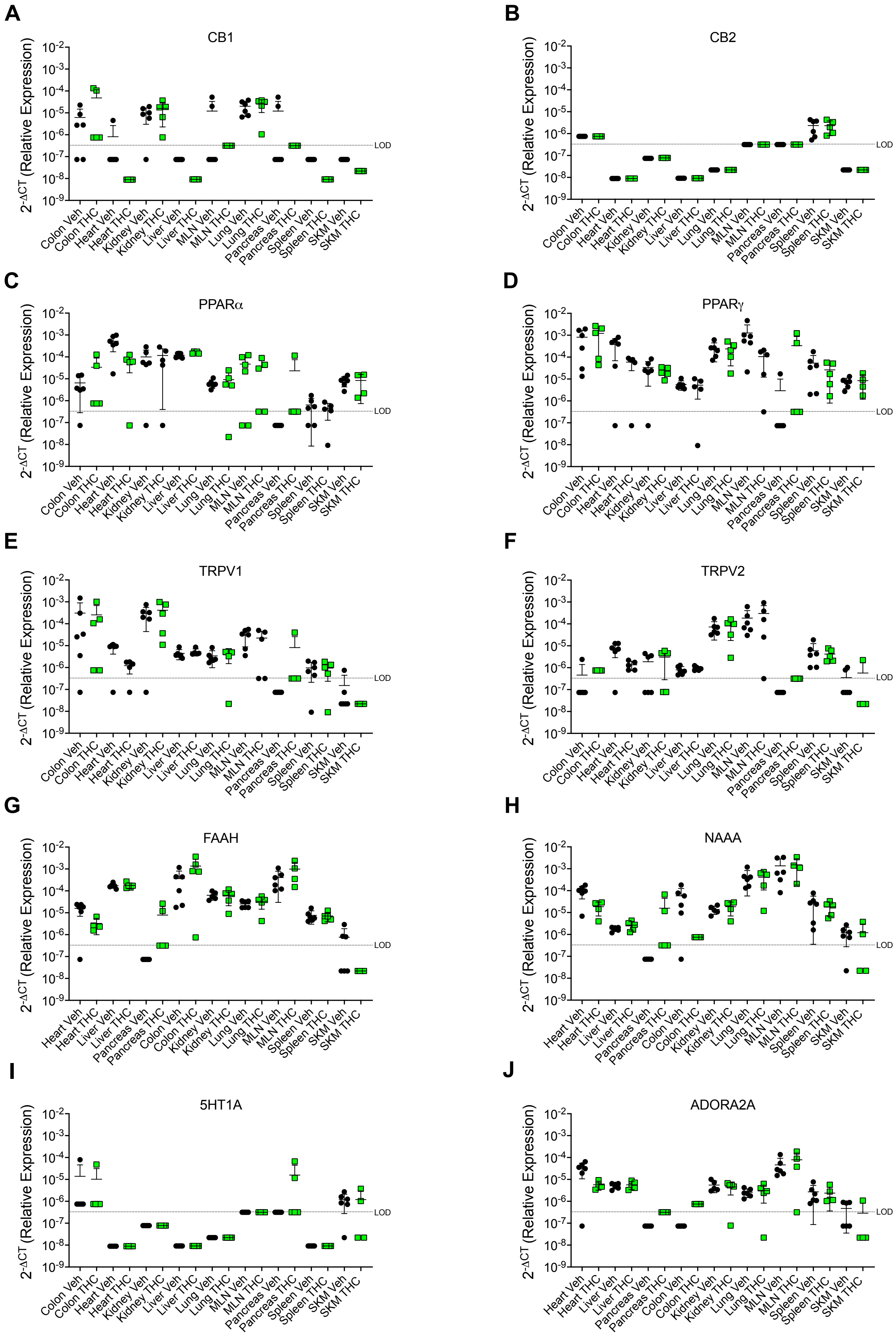

### Supplemental Figure-2

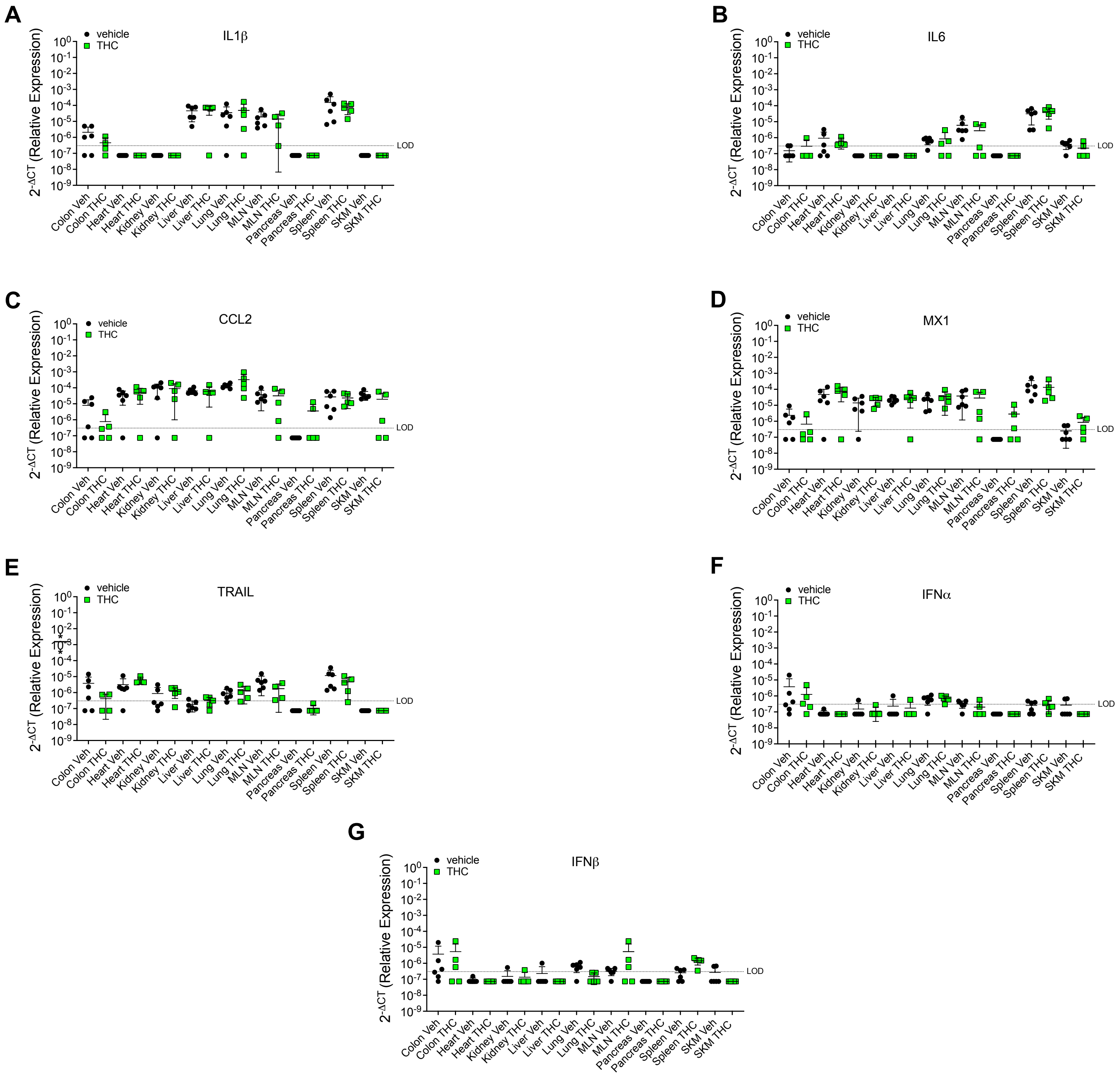
