## Supplemental Table-1 for "Chronic Δ9-Tetrahydrocannabinol Vapor Self-Administration Modulates Gut Microbiota and the Immune and Endocannabinoid Systems in Female Sprague- Dawley Rats"

| Increased Circulating Metabolites | | | |
| --- | --- | --- | --- |
| Metabolite | **Metabolic Pathway** | **Fold Change** | **p-value** |
| N-formylkynurenine | Tryptophan metabolism | 1.62 | 0.0542 |
| Xanthurenate | Tryptophan metabolism | 1.32 | 0.0527 |
| 4-hydroxyphenylacetylglycine | Acetylated peptide | 1.70 | 0.0558 |
| 3-hydroxyoleoylcarnitine | Fatty acid metabolism | 1.92 | 0.0569 |
| Oleoylcholine | Fatty acid metabolism | 2.08 | 0.05 |
| Palmitoyl-linolenoyl-glycerol (16:0/18:3) [2]* | Diacylglycerol | 1.54 | 0.0541 |

| Decreased Circulating Metabolites | | | |
| --- | --- | --- | --- |
| Metabolite | **Metabolic Pathway** | **Fold Change** | **p-value** |
| S-1-pyrroline-5-carboxylate | Glutamate metabolism | 0.41 | 0.0571 |
| Histamine | Histidine metabolism | 0.78 | 0.0548 |
| Serotonin | Tryptophan metabolism | 0.72 | 0.0659 |
| Prolylglycine | Dipeptide | 0.85 | 0.0518 |
| Malonylcarnitine | Fatty acid synthesis | 0.64 | 0.0533 |
